# Pangenome Graph Node-Phenotype Association shows GWAS-like quality results with only few individuals

**DOI:** 10.64898/2026.07.31.741971

**Authors:** Camille Carrette, François Sabot, Cédric Muller

**Affiliations:** DIADE, UM, Cirad, IRD, Av. Agropolis, Montpellier, 34394, France; Genomics Data Analytics, Syngenta Seeds SA, 1228 Chemin de l’Hobit, Saint-Sauveur, 31790, France

**Keywords:** Pangenome, Graph, Association studies, QTL Seq, GWAS

## Abstract

**Purpose:** We introduce GraNPA (Graph Node-Phenotype Association), a method that performs a phenotype-sequence association analysis on a pangenome variation graph (PVG). This PVG is built using a small number of individual genome sequences and does not require additional population materials or kinship information for qualitative phenotypes. This approach minimizes the number of individuals required for association studies and mitigates the impact of reference bias in variant calling within these types of analyses.

**Background:** A PVG is a representation of the multiple alignment of a set of complete genomes. It contains all variations, ranging from single nucleotide polymorphisms (SNPs) to large structural variations (SVs), and these are represented as nodes in the graph. By integrating phenotype information within these nodes, we can assign a Score (*S*) to each one and identify regions related to the phenotype directly within the PVG. These regions represent statistically significant shifts in *S* distribution, indicating their involvement in the phenotype. Finally, GraNPA provides their positions and scores for further analysis.

**Results:** This method was tested using different simulated contexts and two publicly available datasets: the *Sub1A* gene locus for Asian rice *Oryza sativa* in a **13** individuals PVG, and the insertion responsible for the **white-headed** in cattle (*Bos taurus*) with a PVG of **24** individuals. Source code of `GraNPA` is available here https://forge.ird.fr/diade/graphgwas/granpa under GNU GPLv3.

**Conclusion:** `GraNPA` was able to identify the expected area in two simulated datasets, as well as the loci responsible for these two known traits, using only a few dozen complete genomes from these PVGs. Although this method is currently limited to qualitative phenotypes, it paves the way for more efficient methods that rely on PVGs and a small number of individuals.

## 1 Background

### Context

In fields such as plant and animal breeding, physiology, and the etiology of human diseases, identifying the genomic regions responsible for specific phenotypes is often crucial. Genome-wide association studies (GWAS), which use phenotype data and genetic variation in cohorts of hundreds of individuals to identify the genomic regions underlying phenotypes of interest, remain the most popular method to date [1–3]. While the decreasing cost of Next Generation Sequencing has made GWAS more accessible, several critical limitations affect its effectiveness. A fundamental constraint lies in the genotype data acquisition process: indeed, traditional GWAS relies on variant calling through mapping NGS reads to a single reference genome. This approach inherently suffers from reference bias, as potentially missing significant variations present in the population may be absent from the reference genome used [4–7]. Furthermore, the variant calling step and the GWAS analysis itself often incur a substantial computational burden (both in terms of time and resources). Thus, to manage this computational complexity and reduce runtime, GWAS analyses typically rely on pruning methods based on linkage disequilibrium or frequencies to filter out millions of variants. While reducing computational burden, this approach risks the elimination of potentially causal variations from the analysis. Finally, the challenges extend also to the phenotyping, which requires extensive time and human resources to collect data from hundreds of individuals from the genotyped population. This requirement often becomes a bottleneck in GWAS, limiting the scope and scale of analyses. Alternative approaches to GWAS, such as Bulked Segregant Analysis (BSA) and QTL-seq, have also emerged as powerful methods for identifying trait-associated genomic regions. These methods leverage the concept of extreme phenotype sampling by comparing allele frequencies between contrasting phenotypic pools. QTL-seq, for instance, combines BSA with whole-genome sequencing, and offers several advantages over traditional GWAS: it requires fewer individuals, reducing thus phenotyping efforts by focusing on extreme phenotypes, and can be particularly effective in identifying major-effect QTLs. These methods have been successfully applied in various plant species, identifying genomic regions associated with important agronomic traits [8–12]. However, like GWAS, these two approaches still rely on variant calling against a single reference genome, potentially missing important structural variations, and also require at least F2 populations or, even better, Recombinant Inbred Lines (RILs).

### Pangenome variation graphs

As mentioned previously, advancements and cost reductions in second- and third-generation sequencing technologies have transformed genomics, making it possible to generate multiple, high-quality genome assemblies within the same species. This wealth of genomic data and of almost complete genome assemblies provides unprecedented opportunities to capture population-level diversity. While comparing these genomes is crucial for identifying trait-associated genomic regions, managing and analyzing multiple alignments of complex genomes from dozens of individuals presents significant computational and storage challenges. Pangenome Variation Graphs (PVGs) have emerged as an efficient solution to represent and analyze this genomic complexity [13]. These directed graphs provide a compressed yet comprehensive oriented graph structure, where ‘Nodes’ represent shared or unique nucleotide sequences from individual genomes, connected by ‘Edges’ (or Links). Each individual genome can then be reconstructed as a ‘Path’ or ‘Walk’ through the graph — a specific succession of oriented Nodes. This representation, typically stored in Graphical Fragment Assembly (GFA) format, efficiently captures both shared and variable regions across multiple genomes. Several tools have been developed to build PVGs, including VG [14], PGGB [15], and Minigraph-Cactus [16]. While these tools employ different strategies for genome variation identification and PVG construction, and even if the lack of standardization in PVG construction methods remains a challenge in the field [17], the PVGs capture the complete diversity of the individual embedded genomes. This comprehensive representation of the whole catalog of the genomic variations from a population presents an opportunity to develop new approaches for genotype-phenotype association studies.

### GraNPA

To address the limitations of traditional GWAS, QTL-seq, or BSA approaches, while leveraging the comprehensive variation data captured in PVGs, we developed Graph Node-Phenotype Association (`GraNPA`).This novel method adapts the principles of extreme phenotype analysis to PVG structures, combining the statistical power of BSA with the complete representation of genomic diversity offered by PVG, allowing `GraNPA` to overcome several key limitations of existing methods. While traditional approaches require hundreds of individuals and rely on variant calling against a biased single reference genome, `GraNPA` can identify trait association using a dozen individuals by analyzing all genomic variations represented in the PVG. This is achieved by integrating phenotypic information directly with the PVG structure, where the method combines graph topology with phenotyping data to assign scores to individual nodes, thereby identifying genomic regions associated with traits of interest. The effectiveness of `GraNPA` has been validated using simulated data and two published datasets (animal and plant), demonstrating its ability to identify trait associations previously found through traditional methods, but with much less individuals and a higher precision. The method requires minimal inputs: a single PVG in GFA v1.1 format and the phenotype information for some or all individuals represented in, with at least 3 contrasting individuals. The output maintains familiarity with traditional approaches by generating GWAS-style visualisations, including Manhattan plots, while offering enhanced resolution by providing both genomic ranges and specific graph nodes associated with traits of interest. GraNPA is a member of the GraSuite (https://forge.ird.fr/diade/GraSuite) and is available as a GNU GPL3 code with an associated Singularity container at https://forge.ird.fr/diade/graphgwas/granpa.

## 2 Implementation

### 2.1 GraNPA pipeline Methodology

As shown in Figure 1, in the first step of the pipeline, the PVG, in GFA v1.1 format (with genomic start and stop positions of each walk), is indexed by GraTools [18], which the user must install (included in the Apptainer), to identify which individuals each node belongs to. Using this information, stored in the BAM file generated by GraTools, alongside the phenotype data, a phenotypic score is computed for each node, as in the Figure 2: each time an individual with a given phenotype will go through the node, it will modify the node score by its own value (-1 if negative phenotype, +0 if unknown, and +1 if positive). Then, the *rank compare 2indep* function (from package *statsmodels*, [19]) is applied to a range of consecutive nodes to detect changes in the score distribution. The *p*-values from this test are displayed in a Manhattan plot, and a summary of the 20 windows with the lowest *p*-values is outputted with information about the nodes (positions from the GraTools bed files). This pipeline has been validated with different contexts of simulated data and with two use cases with known trait location: the Asian rice *Oryza sativa* submergence resistance [20], supported by a graph of 13 individuals, and the cattle *Bos taurus* white-head [21] trait, with a graph of 24 individuals.

**Fig. 1.**
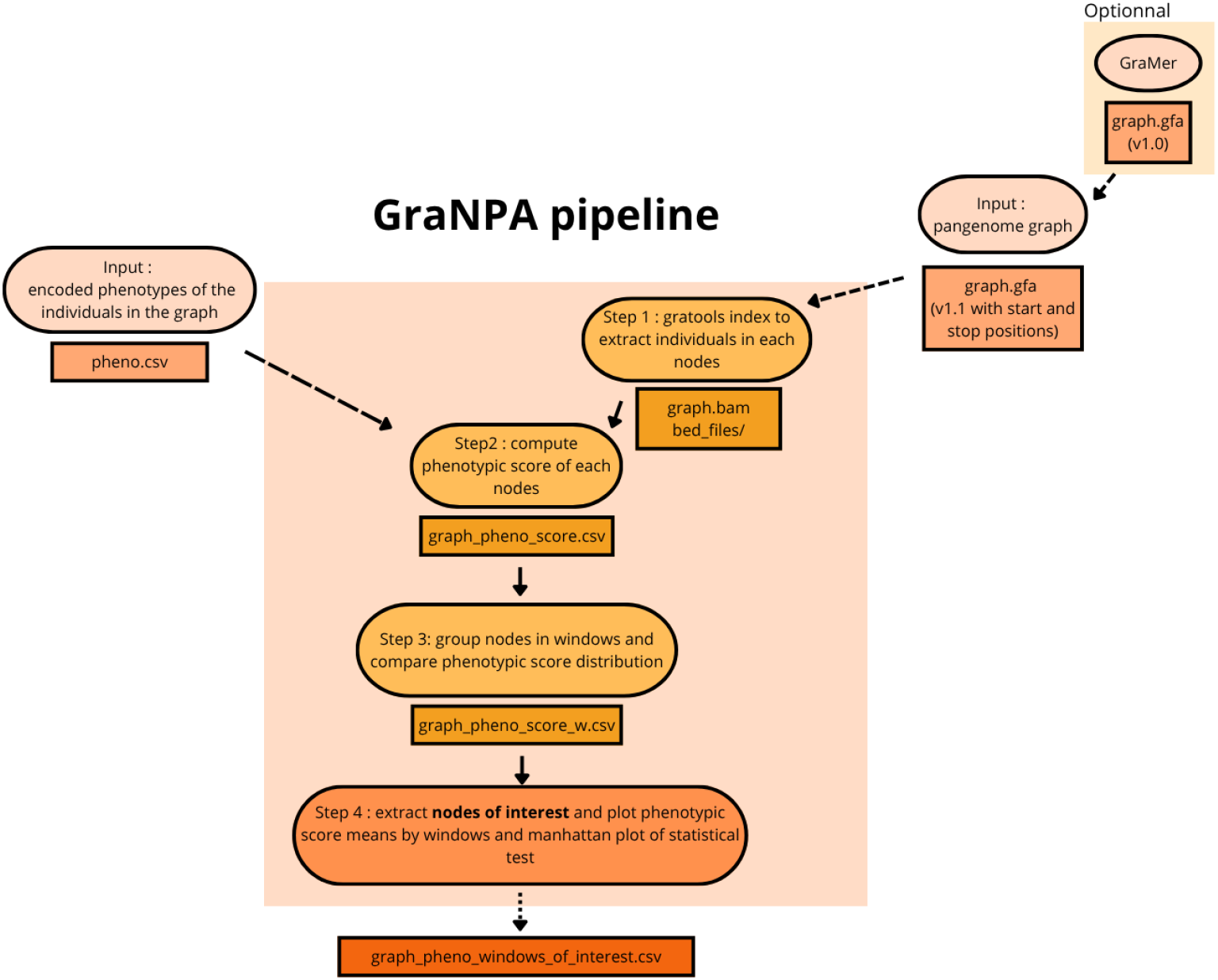
GraNPA pipeline overview. `GraNPA` takes as input a PVG in GFA v1.1 format (with the genomic start and stop positions of walks), or from a GFA v1.0 converted with `GraMer`, and an encoded phenotype values table (+1, 0, or -1) for each individual/haplotype of the PVG in a CSV file. The pipeline first indexes individuals present at each node using GraTools [18], then computes node-level phenotypic scores. Nodes are grouped into consecutive windows, and phenotypic score distributions are compared between groups using the *rank compare 2indep* statistical tests. Intermediate results are stored as temporary CSV files, allowing users to re-run analyses with different window sizes or visualization options without recomputing scores. The main output is a ranked list of candidate windows sorted by ascending *p*-value (most significant first), with their mean scores, *p*-values, and chromosomal locations. Optional visualization outputs include scatter plots of window means and Manhattan-like plots of *p*-values across chromosomes.

**Fig. 2.**
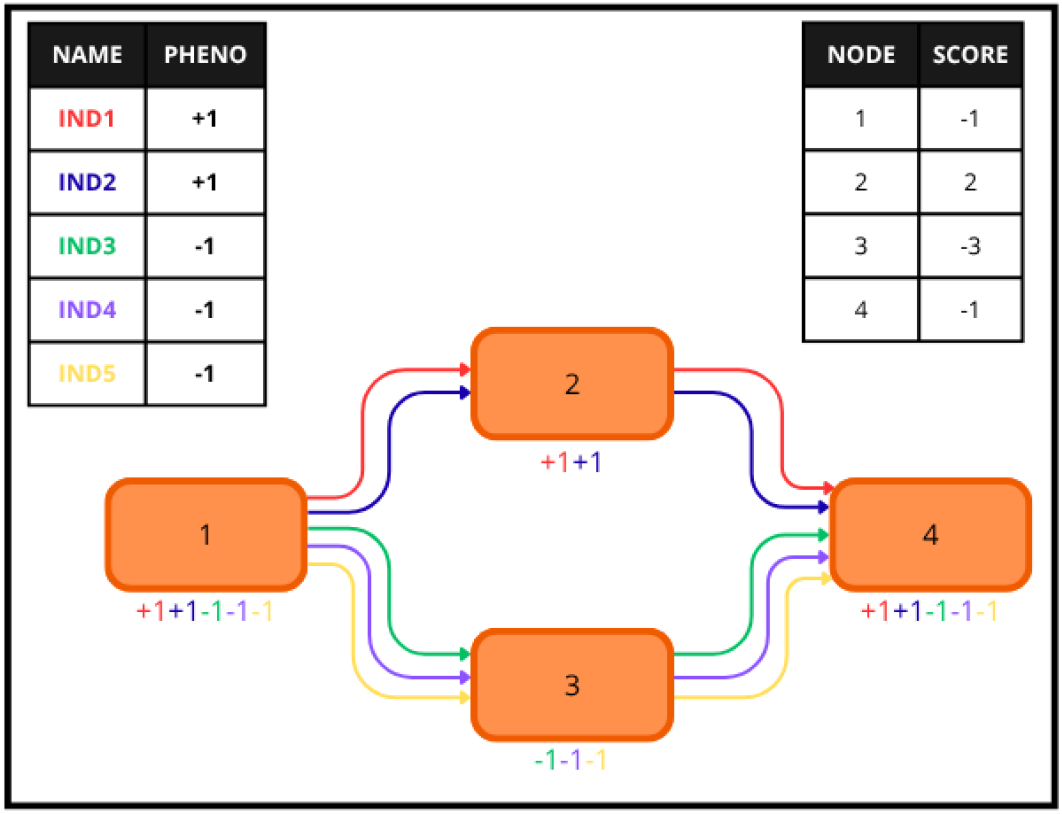
Step 2 of the GraNPA pipeline. The top-left table contains the expected encoded phenotype for each individual of the graph as a CSV file (input file). Individual phenotypes are encoded +1 when carrying the trait of interest, -1 when not carrying and +0 when the phenotype is unknown or not sure. The middle structure is a representation of the graph with the Walk of each individual. Each individual which crosses a node based, on their Walk, will add their encoded phenotype to the node. A final score is calculated by summing the encoded phenotype for each node and the result is shown in the top-right table. Node 2 and node 3 present respectively the maximum positive score (*S*_*max*_) and the maximum negative score (*S*_*min*_) that are the 2 extreme scores that the nodes of the graph could have.

### 2.2 PVG simulation

The simulated graph was created by generating a random DNA sequence (G0) of 170 kb, composed of two chromosomes (100 kb and 70 kb). Random variations (indel rate of 0.005 with a maximum size of 50 bp and SNPs at a 0.01 rate) were then applied ten times separately to G0 to create G1 to G10. Each of these ten genomes was then used to generate two additional genomes, to whom the same random variation script was applied, generating G1 1, G1 2, G2 1, G2 2, and so on (see Appendix A.1). G0 was subsequently removed to avoid including the “common ancestor” in the graph, leaving a dataset with genomes which shared variations but also had specific ones. Before constructing the graph, variations specific to certain simulated individuals had to be created. Seven scenarios were created to test `GraNPA` on a wide range of possibilities. The most common scenarios are the deletion and insertion cases, which are described below. The other five follow the same generation method with specific changes to the genomes, which are described in A.2-A.6. For the insertion case, a 5 kb segment (positions 45–50 kb of chromosome 1) was deleted from 24 genomes, so that only the six remaining genomes (G3, G3 1, G3 2, G7, G7 1, and G7 2) retained this segment. This shared, specific presence defined the positive phenotype in the `GraNPA` run. Conversely, for the deletion case, this 5 kb segment was removed exclusively from these same six positive-phenotype individuals, while the remaining 24 genomes retained it. The 30 genomes were then used to construct a PVG with Minigraph-Cactus in full mode. All codes and command lines used for building these PVGs are available at [the GraNPA DataSuds repository](https://dataverse.ird.fr/dataverse/granpa).

### 2.3 PVG origin

The rice pangenome PVG was constructed using Minigraph-Cactus with the 13 Asian rice *O. sativa* individuals from [22], as described in [23], and is already compatible with `GraNPA`. The cattle *B. taurus* PVG was retrieved from https://zenodo.org/records/15097551 and was built from 24 publicly available individuals [21] using PGGB, resulting in 29 GFA files (one GFA per chromosome). Since PGGB outputs multiple GFA v1.0 files, these GFA files were processed using `GraMer` (https://forge.ird.fr/diade/graphgwas/gramer). As all GFAs from PGGB have nodes with IDs beginning with the number 1 and continuing to the last node, merging those GFAs would cause confusion regarding the IDs of the nodes. To ensure the IDs are unique, the last ID of chromosome 1 is added to the IDs of chromosome 2. This process was repeated for each subsequent chromosome. Thus, GraMer converted the PGGB outputs into a single GFA v1.1, e.g. specifying the genomic start and stop positions of the walks.

### 2.4 Phenotypic score calculation and filtering

Node scoring was performed using GraTools [18] *import* command to identify information about individual Walks through the PVG. This scoring process requires theidentifiers of individuals in the input phenotype file to match those in the PVG. A rein-forcement algorithm was implemented where individuals exhibiting the trait of interest were assigned a weight of +1, those lacking the trait were assigned − 1, and individuals with missing phenotype data were assigned +0. For each node, the algorithm computes a cumulative score *S* computed by summing the weights of all individuals traversing it (Figure 2). The resulting scores range from a theoretical minimum *S*_*min*_ (nodes traversed exclusively by individuals lacking the trait) to a maximum *S*_*max*_ (nodes traversed exclusively by individuals carrying the trait). To minimize topological noise and enhance the detection of truly discriminative nodes, we implemented a simple adaptive filtering strategy. A dynamic threshold *θ* was defined as:

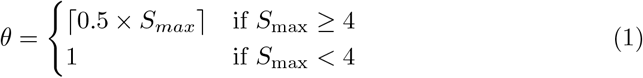

All nodes with a score *s* such that *S*_*min*_ *< s < θ* were reset to zero. This approach keeps the *S*_*min*_ and 60% of the highest positive scores while removing noise. The threshold is set to 1 if the maximum number of individuals with a positive phenotype in the graph is lower than 4. This was set up in order to keep more information if the *S*_*max*_ is low. For instance with the cattle data (4 individuals carrying the trait out of 24), *S*_*min*_ = 20, *S*_*max*_ = 4, and *θ* = 1.6; all nodes with a score strictly between 20 and 1.6 are set to zero, retaining only scores of 20, 0, 2, 3, and 4 for the analysis.

This filtering step partitions the nodes into three functional categories:

- **Nodes with a score of zero:** Considered uninformative but retained in the analysis to provide structural context.
- **Nodes with high positive scores:** Found in at least 60% of trait-carrying individuals, representing candidate nodes for causal insertions.
- **Nodes with the minimum phenotype score (***S*_*min*_): Present exclusively in non-trait-carrying individuals, representing candidate nodes for causal deletions.

### 2.5 Phenotypic scores comparison

To identify genomic regions associated with the trait of interest, nodes are grouped into consecutive windows (default: 400 nodes with an overlap of half the window size; user-configurable) based on their topological IDs generated by the graph constructor. Windows are constructed independently for each chromosome.

For each window, two complementary metrics are computed: (*i*) the **Mean phenotypic score:**, that indicates the average strength of trait association within the window, and (*ii*) the ***p*-value:** that quantifies whether the window’s score distribution differs significantly from the chromosome-wide score distribution.

We compared scores within each window (*X*_1_) to scores in the chromosome background (*X*_2_) using the *rank-based two-sample* procedure (*rank compare 2indep* function) from the Python *statsmodels* package [19]. The method estimates the probabilistic index (relative effect) as following:

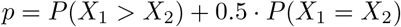

and tests (*H*_0_: *p* = 0.5) against (*H*_*A*_: *p >* 0.5) (window stochastically larger than background). The *p*-values for this test are derived from a normal approximation for large samples, which handles extensive ties (repeated values) more effectively than the classic Mann–Whitney test. Windows with the lower *p*-values and high mean phenotypic scores indicate regions enriched for nodes associated with the trait.

### 2.6 Output

Finally, `GraNPA` provides various outputs so the user can interpret results. Firstly and by default a CSV with the top 20 windows of interest sorted by their respective *p*-value. For top-ranked windows, detailed information is provided, including their genomic coordinates mapped to linear genomes for individuals carrying those nodes in their paths, the node-level phenotypic scores, and the local visualization plots. A filter is applied to these windows, only those carried by all or none of the individuals with the trait are considered candidates. Genome-wide Manhattan plots are also generated, displaying *p*-values and mean scores across all chromosomes, with segments ordered by topological IDs along the x-axis.

## 3 Results

### 3.1 Simulated data

In order to demonstrate the reliability and the limits of the `GraNPA` pipeline, it was applied to seven simulated scenarios dataset that was generated (see 2.2). `GraNPA` was tested on two scenarios that are explained in the text and five more are available in Appendix A.2 to A.6. Results from insertion and deletion cases was made by running `GraNPA` on two PVG built from 30 artificial genomes spanning 170kb upon 2 chromosomes (see section 2): one was constructed to integrate a 5kb insertion to 6 genomes, and the second a deletion of 5kb in 6 genomes also. The expected result location is 45,000–50,000bp in both cases. `GraNPA` was run following the pipeline describe in 2.1, the *p*-value output are shown in Figures 3 and 4. The windows highlighted by `GraNPA` are overlapping the expected results for both cases. It is quite important to note that, in the deletion test, as expected, these windows do not contain positions on the genomes of the individuals of interest (as they do not harbor the deleted nodes), and that the average phenotypic score for these windows is negative, as expected (see Supplementary table A.1).

**Fig. 3.**
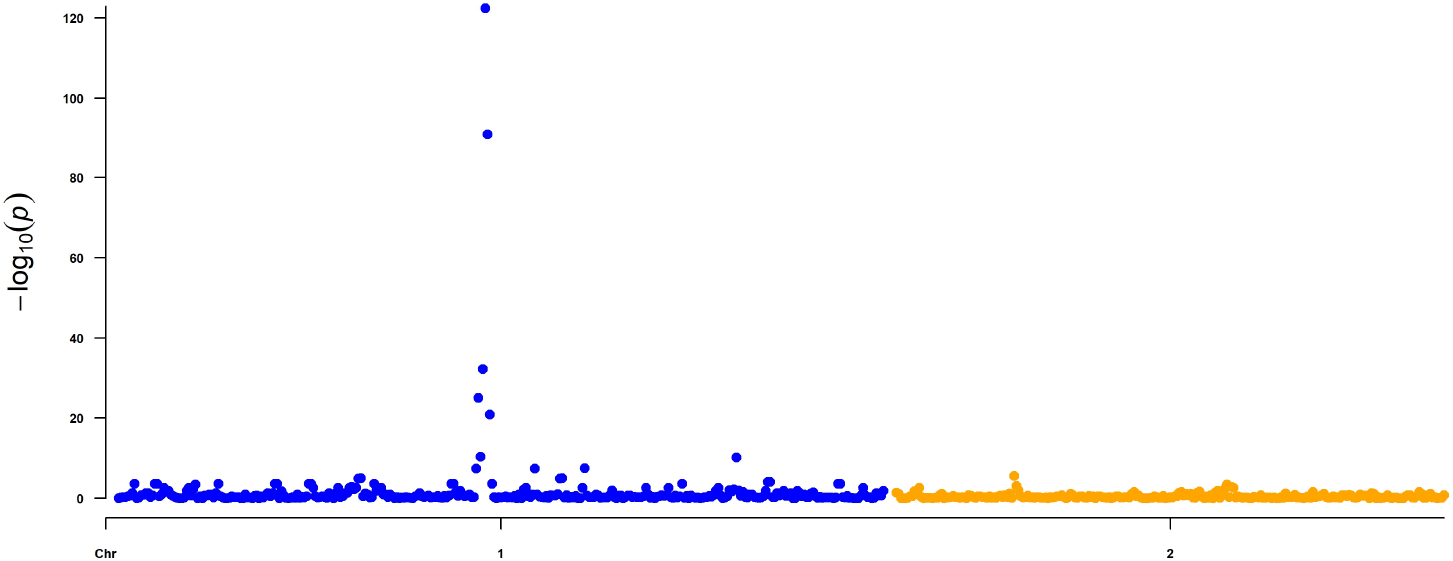
*P*-values plot of *rank compare 2indep* test on 400 node windows on a simulated PVG of 30 individuals with a simulated causal insertion for 6 samples. The x-axis shows the ID of the first node of each window analyzed across the (pan-)chromosomes (in blue the 100kb and in yellow the 70kb) in the PVG, and the y-axis shows the -log of the *p*-values. Peaks in the *p*-values show a significant change in the distribution of the phenotypic score.

**Fig. 4.**
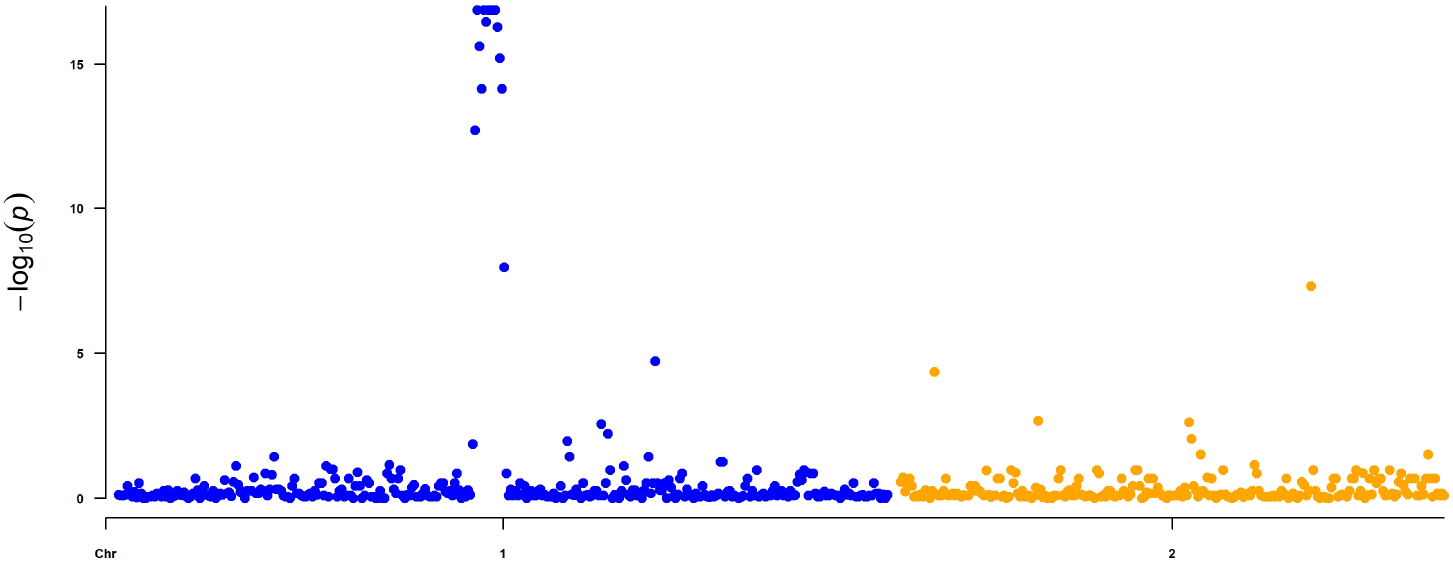
*P*-values plot of *rank compare 2indep* test on 400 node windows on a simulated PVG of 30 individuals with a simulated causal deletion for 6 samples. The x-axis shows the ID of the first node of each window analyzed across the (pan-)chromosomes (in blue the 100kb and in yellow the 70kb) in the PVG, and the y-axis shows the -log of the *p*-values. Peaks in the *p*-values show a significant change in the distribution of the phenotypic score.

### 3.2 Asian rice *Oryza sativa* Sub1A gene

#### Expected results

Submergence resistance in Asian rice *O. sativa* is linked to a gene family comprising three genes: *Sub1A, Sub1B*, and *Sub1C*. Among these, only the *Sub1A-1* allele confers submergence tolerance [20]. We used here the 13 *Oryza sativa* genomes PVG created in [23] and made from the Platinum rice genomes [22]. The complete PVG consists of 26,461,214 nodes with a mean length of 32 bp (excluding SNPs), while the top 5% longest nodes have an average length of 508 bp [18, 23].

While all 13 lines harbor *Sub1B* and *Sub1C* genes, only four of the *indica* lines carry the *Sub1A* gene and only two carry the *Sub1A-1* allele of the gene giving the sub-mergence resistance. `GraNPA` was run with the submergence phenotype and expected locus were found but drawn with other results (rank position 121 and 221) certainly due to the low number of individual carrying the submergence trait (see Appendix Figure A.7). The submergence genotype were then converted in a presence/absence of *Sub1A* where the sub1A-1 or Sub1A-2 isoform are located in a insertion carried by four of the individuals of this dataset and this SV being absent from all other lines including the *japonica* Nipponbare reference genome (see Appendix Table A.2). This change was made to test the hypothesis of the dilution of the results due to the number of individuals carrying the trait. We tried to localize the *Sub1A* gene using only the 13 lines of the graph, without additional genomic material. We thus encoded the presence of *Sub1A* as a binary phenotype, assigning +1 to the four *indica* lines harboring the gene (whatever its allele), and -1 to the remaining nine accessions.

We used the reported genomic position of *Sub1A* from the literature [24] (IR64 genome, chromosome 9: 7,546,385–7,547,220) as control. As `GraNPA` operates on the PVG nodes rather than on the genomic coordinates, identifying the nodes corresponding to *Sub1A* provides a ground truth for validation. Using GrAnnoT [23], we mapped the IR64 *Sub1A-2* copy to the following graph nodes: 25,250,739; 25,250,741; 25,250,743; 25,250,744; 25,250,746; and 25,250,747. Two additional nodes, 25,250,742 and 25,250,745, represent the alternate alleles at the SNP sites corresponding to nodes 25,250,743 and 25,250,746, respectively, and distinguish the *Sub1A-1* and *Sub1A-2* isoforms.

#### Identification of the locus on Chromosome 9

The `GraNPA` pipeline, as described in Section 2.1, was applied to this rice dataset. The total execution time, including the `GraTools` preprocessing step, was approximately of 2 hours using 16 CPU cores (with an additional 4h for computing the PVG [23]). The resulting *p*-value distribution is shown in Figure 5. In this context, the *p*-value represents the probability that the phenotypic score distribution within a given window deviates significantly from the overall chromosomal distribution randomly. The rank-based distributional analysis identified two major, significant peaks across the genome on chromosomes 11 and 9 (see figure 5). After selecting windows that are present in all the four (or none) of *sub1A*-carrying individuals, the top two windows overlap the known location of the gene on chromosome 9 at the expected location of the *Sub1A* gene, spanning respectively from nodes 25,250,429 to 25,250,829 and from 25,250,629 to 25,251,029. These overlapping windows correspond to coordinates chr9:7,528,190-7,552,127 and chr9:7,540,994-7,562,784 on the IR64 accession (see table 1), and are represented in red on Figure 5.

**Table 1.** Summary of the most significant windows identified by `GraNPA` on the *Oryza sativa* PVG and having positions on the individuals that carry the trait. Overlapping windows consist of 400 consecutive node IDs and are labeled by their start and end node IDs, and the (pan-)chromosome on which they are located. The mean phenotypic score and the *−*log_10_(*p*-value) for each window are provided. Finally the genomic coordinates on Os117425RS1 (IR64 genome) are displayed. The two highlighted windows (in bold) overlap with the expected *Sub1A* locus linked to the submergence tolerance trait.

| **window_id** | **node_start** | **node_stop** | **chr** | **mean** | **pvalue** | **OsIR64RS1** |
|---|---|---|---|---|---|---|
| **1** | **25,250,629** | **25,251,029** | **9** | **2.485** | **213.72** | **chr9:7540994-7562784** |
| **2** | **25,250,429** | **25,250,829** | **9** | **2.4025** | **189.43** | **chr9:7528190-7552127** |
| 3 | 25,250,829 | 25,251,229 | 9 | 2.13 | 128.71 | chr9:7552033-7574390 |
| 4 | 5,899,098 | 5,899,498 | 11 | 1.725 | 114.3 | chr11:19080979-19096803 |
| 5 | 21,956,076 | 21,956,476 | 7 | 1.8025 | 113.59 | chr7:24062666-24076830 |
| 6 | 8,254,906 | 8,255,306 | 12 | 1.64 | 108.84 | chr12:12318990-12324504 |
| 7 | 25,257,429 | 25,257,829 | 9 | 2.3925 | 106.6 | chr9:7587559-7621155 |
| 8 | 4,359,621 | 4,360,021 | 10 | 2.06 | 104.66 | chr10:3814746-3841395 |
| 9 | 24,560,630 | 24,561,030 | 8 | 1.69 | 97.5 | chr8:27898329-27900514 |
| 10 | 21,956,276 | 21,956,676 | 7 | 1.655 | 96.84 | chr7:24062666-24076834 |

**Fig. 5.**
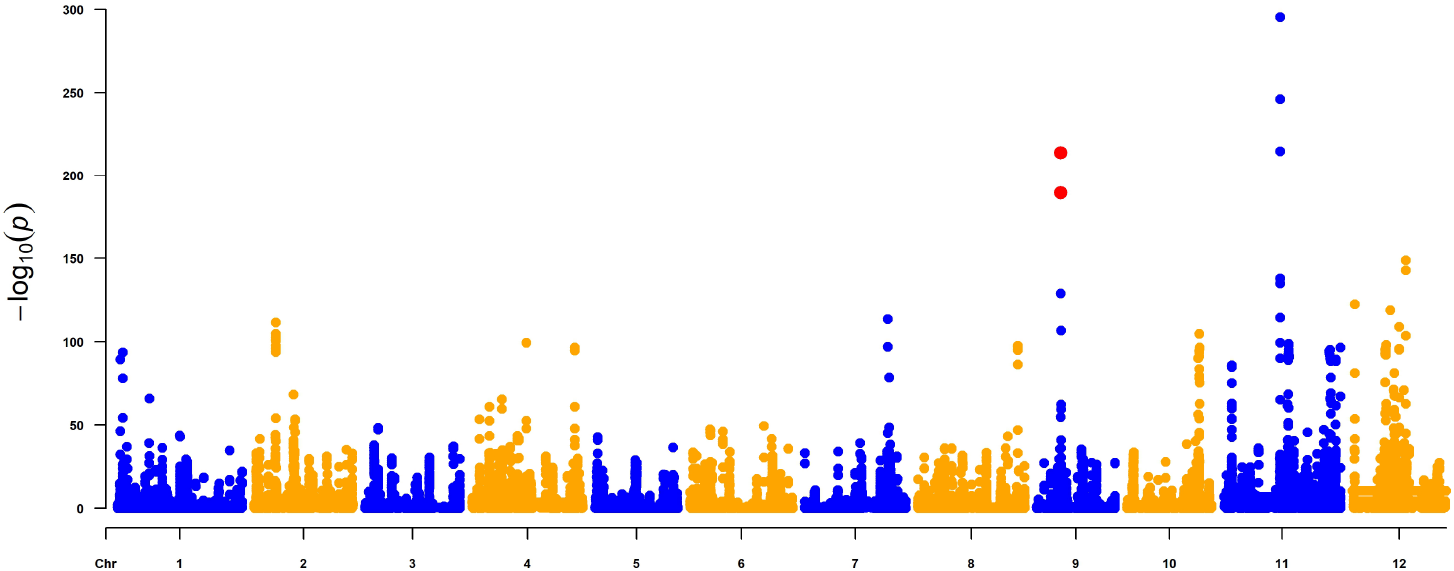
*P*-values plot of *rank compare 2indep* test on windows of 400 nodes on *Oryza sativa* PVG with the highlight (red dot) of the windows overlapping the *Sub1A* known position. The y-axis shows the -log of the *p*-values, and the x-axis shows the ID of the first node of each window analyzed across the (pan-)chromosomes in the PVG. The alternating colors indicate the change in chromosomes to make the plot easier to read. Peaks in the *p*-values show a significant change in the distribution of the phenotypic score.

### 3.3 Cattle White-head

#### Expected results

A cattle (*Bos taurus*) PVG comprising 24 long-read genome assemblies was built using PGGB, alongside 250 short-read samples, to identify the causal variationresponsible for the white-headed phenotype [21]. Among the 24 individuals embedded in this PVG, four (one Holstein and three Simmentals) exhibit the white-headed trait (see Appendix Table A.3). The phenotype was encoded as +1 for the individuals with the white-head trait, and -1 for the rest. In the Holstein reference genome, the locus associated with this phenotype is an insertion on chromosome 6 (positions 70,099,532–70,120,129), which corresponds to nodes 14,192,067 to 14,204,232 in the PVG. `GraNPA` performed an association analysis across all the 29 chromosomes to evaluate its ability to recover this known variation.

#### Preprocessing

An additional preprocessing step was required because the PGGB output format is not natively compatible with the GraTools pipeline. PGGB generates separate GFA v1.0 files for each chromosome, which results in two main issues: first, node IDs are not unique across the different files, whereas `GraNPA` requires a global indexing system; second, PGGB utilizes P lines (Path), which describe node orientations using “+” or “-” signs and lack explicit genomic coordinates, while our pipeline requires the `W` (Walk) format.

To address these formatting challenges, we used `GraMer` (https://forge.ird.fr/diade/graphgwas/gramer), a custom Python script designed to merge individual chromosome GFAs into a single PVG file with a unified indexing system. The GraMer run took about 13 minutes with 16 CPU cores, resulting in a merged GFA v1.1 file (13 GB) comprising 52 million segments, each averaging 70 bp in length.

#### Identification of the locus on Chromosome 6

The analysis performed by `GraNPA` across the entire graph (comprising 52 million segments, analyzed using overlapping 400-node sliding windows) took approximately 5 hours using 16 CPU cores (excluding the PVG creation and its conversion time).

Figure 6 shows the identification of 5 main peaks (chromosomes 4, 6, 12, 15 and 22). After applying a filter based on ownership of the windows (retaining only those belonging to all or none of the individuals of interest), the 20 first better windows contain two adjacent windows with a high statistical significance in immediate proximity to the 20 kb insertion previously described by [21] (expected positions: chromosome 6: 70,099,532–70,120,129 on the reference genome; nodes 14,192,067–14,204,232).

**Fig. 6.**
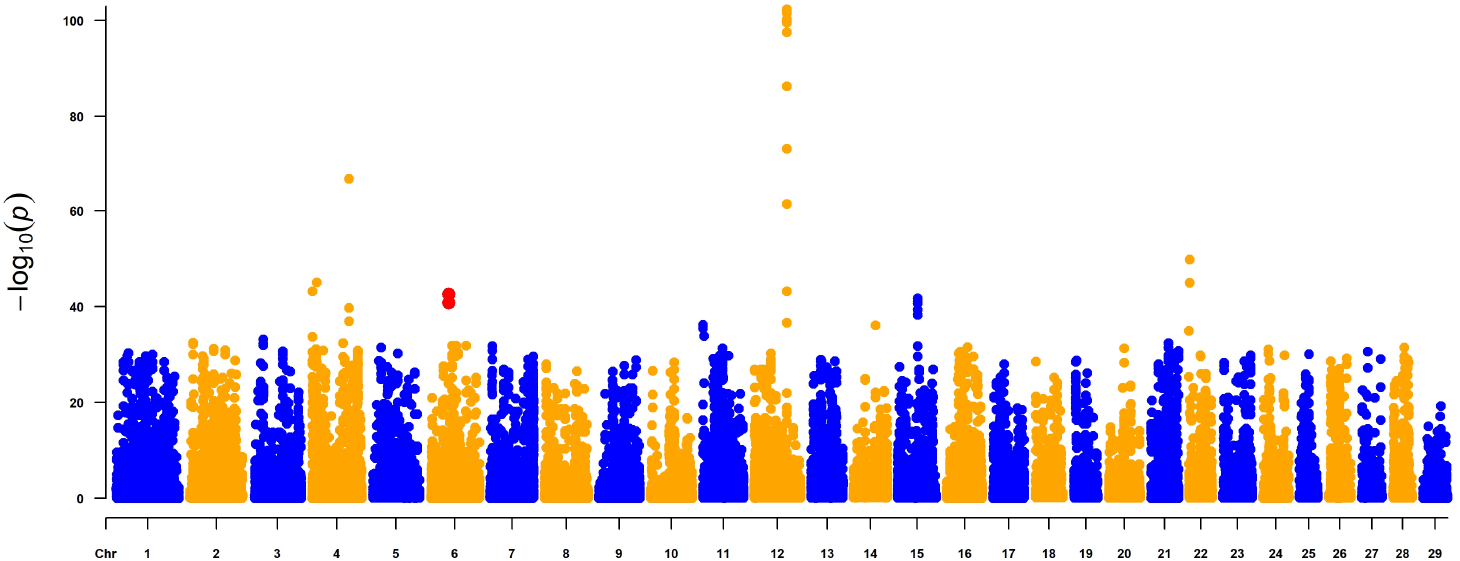
*P*-values plot of *rank compare 2indep* test on windows of 400 nodes on the cattle PVG. The y-axis shows the -log of the *p*-values, and the x-axis shows the ID of the first node of each window analyzed across the chromosomes of the PVG. Peaks in the *p*-values show a significant change in the distribution of the phenotypic score. The alternating colors indicate the change in chromosomes to make the plot easier to read.

These two windows, which ranked 7^th^ and 8^th^ with *p*-values of − log_10_(*P*) = 42.55 and (− log_10_(*P*) = 40.79) span respectively from nodes 6: 14,204,077 to 14,204,477 and from nodes 14,203,877–14,204,277, corresponding to positions 70,103,306-71,037,322 on the Holstein reference genome (see Table 2).

**Table 2.** Summary of the most significant windows identified by GraNPA on cattle and having positions on the individuals that carry the trait. Windows of 400 successive node IDs are described by their start and end node IDs, and the (pan-)chromosome where they are located. The mean phenotypic score and the *−* log_10_(*p*-value) for each window are provided. Finally the genomic coordinates on the ARS UCD12 Holstein reference are displayed. The two highlighted windows (in bold) overlap with the expected insertion linked to white head trait.

| **Window ID** | **Start Node** | **End Node** | **Chr** | **Mean** | **-log(p-value)** | **ARS_UCD12** |
|---|---|---|---|---|---|---|
| 1 | 28169845 | 28170245 | 12 | 1.6125 | 86.24 | chr12:70091482-70109205 |
| 2 | 28171445 | 28171845 | 12 | 1.4550 | 73.20 | chr12:69904887-70076405 |
| 3 | 10225156 | 10225556 | 4 | 1.0800 | 66.67 | chr4:94294710-94308667 |
| 4 | 28170245 | 28170645 | 12 | 1.3275 | 61.41 | chr12:70076405-70091483 |
| 5 | 8598356 | 8598756 | 4 | 0.8550 | 45.05 | chr4:8229752-8236326 |
| 6 | 28169645 | 28170045 | 12 | 1.0150 | 43.18 | chr12:70105913-70111199 |
| **7** | **14204077** | **14204477** | **6** | **1.2625** | **42.55** | **chr6:70103306-71037322** |
| **8** | **14203877** | **14204277** | **6** | **1.2175** | **40.79** | **chr6:70103306-71011975** |
| 9 | 10224956 | 10225356 | 4 | 0.7450 | 39.69 | chr4:94294710-94298275 |
| 10 | 10225356 | 10225756 | 4 | 0.7050 | 36.96 | chr4:94294852-94308719 |

## 4 Discussion

### Simulated dataset

As expected, `GraNPA` highlights nodes specific to individuals with a requested phenotype, yielding the lowest *p*-values in both insertion and deletion cases. For other genomic cases (available in appendix), `GraNPA` was able to identify a simulated area linked to a phenotype in conditions similar to the rice and cattle datasets (see Appendix panels A.2 and A.3). We also showed that a phenotype linked to two different regions in the genome could be retrieved by the method (see Appendix A.5). In order to have a negative control, we also run `GraNPA` on a dataset with randomized phenotypes, and the tool did not identify any peaks in the graph (see Appendix A.6. Finally, we ran the pipeline on a phenotype linked to a unique single simulated SNP. Unfortunately `GraNPA` was not able to retrieve the expected area (see Appendix A.4). This result shows a limit of our method: `GraNPA` relies on a minimum number of nodes that is associated with the trait. A unique single SNP is not enough to change the 400 nodes window score compared to other random chromosome windows. However, this extreme scenario may be uncommon in empirical datasets. Causal variants are often in linkage disequilibrium with neighboring polymorphisms, resulting in multiple correlated variants-and therefore multiple graph nodes-associated with the phenotype, even when only one variant is causal. This local enrichment is essential for `GraNPA`: when few or no phenotype-associated nodes are shared among individuals in the vicinity of a causal locus, the method is expected to have limited power to pinpoint the region.

These simulations demonstrate the ability of `GraNPA` to detect the specificity of a given binary phenotype across the PVG with few individuals. Indeed, `GraNPA` does not test direct association, but rather uses the PVG to propagate the phenotypic score and observe the behavior of these data across the nodes. Furthermore, the simulation highlights that the *p*-value is not the sole critical output of `GraNPA`; the mean phenotypic score and the position of the windows on the individuals of interest can also guide the interpretation of the results.

### Effectiveness with Small Sample Sizes

The primary objective of our study is to show the ability of `GraNPA` to identify causal loci using a remarkably limited cohort. In the Asian rice case study, the *Sub1A* locus was found using only 13 individuals. Similarly, the cattle White Head trait was pin-pointed with a cohort of only 24 genomes, while the original study needed the graph and an additional 250 short-read sequencing samples [21]. Traditional GWAS typically require hundreds to thousands of samples to reach sufficient statistical power. By leveraging the structural information inherent in PVGs, `GraNPA` acts as a “prioritization compass”, drastically reducing the search space to a few candidate windows, even when biological material is scarce. As shown in the *Sub1A* analysis (Figure 5), only four sets of windows across the entire genome reached such high significance levels, meaning that an investigator without prior knowledge of the locus would only have to inspect only four limited genomic regions. Despite the limited sample size of 13 assembled rice genomes and of 24 cattle genomes, the method achieved in both cases a substantial reduction in search space, isolating the causal locus among the (maximum 10) top-ranking signals, genome-wide with each region about respectively 13 Kbases (400 nodes x 32 bases mean length) in rice and about 28 Kbases for bovin (400 x 70 Kbases mean length).

This behaviour contrasts with conventional GWAS, in which variants are typically tested independently and statistical power depends strongly on the number of sampled individuals available to support each marker-level association. In `GraNPA`, association is assessed at the level of genomic windows rather than individual variants. For each chromosome, the distribution of phenotypic scores within a window is compared with the chromosome-wide background distribution. Consequently, a genomic region can be detected when it contains a local enrichment of nodes with high phenotypic scores, even when the number of individuals is limited. The statistical evidence therefore derives from the aggregation of concordant node-level signals within a window and from their contrast with the chromosomal background, rather than from independent tests of individual variants alone. However, this advantage depends on a sufficiently strong and shared local signal: if too few genomes support the phenotype, or if their phenotypic signal is weakened by label noise, the relevant window may no longer be distinguishable from the chromosomal background.

We therefore examined the lower limit of `GraNPA` under conditions of extreme class imbalance and phenotype-label perturbation. In rice, the submergence-tolerant *Sub1A-1* isoform was present in only 2 of the 13 genomes (with 2 genomes carrying *Sub1A-2* and 9 genomes lacking both isoforms). Under this setting, the expected *Sub1A-1* signal was no longer clearly separated from background and became obscured by noisy windows more associated to population structure rather than the trait itself (see Appendix A.7). In cattle, where only 4 of 24 genomes displayed the White Head phenotype, we simulated phenotype noise by masking two positive genomes (phenotype +1 → 0). The target region remained detectable after filtering, but its rank dropped from 7–8 to the end of the top 20, and its significance decreased, bringing it closer to the reporting cut-off. Taken together, these stress tests indicate that, while `GraNPA` can operate with small cohorts, its performance degrades rapidly when fewer than three genomes support the phenotype; we therefore recommend at least three phenotype-supporting individuals, and cautiousness when interpreting results below this threshold.

### Signal Ranking

Genome-wide analysis of the cattle dataset (Figure 6) identified approximately six distinct peaks. While the Manhattan representation effectively visualizes the spatial distribution of phenotypic scores across the PVG, *p*-values alone — representing statistical distribution comparisons — are insufficient for a comprehensive assessment. To refine this signal, we implemented a stringent quality filter, retaining only windows with balanced presence/absence patterns. This approach consistently ranked the expected region on chromosome 6 within the top 8 most significant windows out of 52 million analyzed segments. This high-ranking performance, achieved despite the high dimensionality of the graph, validates the robustness of our ranking methodology in distinguishing biological signals from topological noise.

This topological noise found in the cattle results however highlighted an important aspect of the PVG approach in `GraNPA`: the process expects the PVG to be perfect with perfect assemblies, which we know it is not yet the case. Indeed, most of the non-specific peaks we observed such as the peaks in the chromosome 4 and chromosome 11 on Figure 6 are linked to telomeric regions, which are known to be complex to assemble and to include in the PVG with the current version of the different tools. However, it allows the results to improve with better PVGs in the near future.

### Advantages of a Topology-Only Approach

Operating exclusively on the PVG topology offers a low-cost and bias-free alternative for phenotype inspection. Unlike existing methods that rely on mapping short reads to a PVG and coming back to a reference-based VCF to perform a classic GWAS [25], `GraNPA` eliminates the reference bias and the heavy computational burden of alignment. Furthermore, the use of phenotype scores provides a direct interpretation of the variant type: a high maximum phenotype score identifies nodes present exclusively in individuals carrying the trait, suggesting a causal variation (SNP or insertion). Conversely, a high minimum phenotype score identifies nodes present only in individuals lacking the trait, suggesting also a causal variation (probably a deletion). This dual-signal approach allows for a more nuanced understanding of how SVs drive phenotypic diversity. In addition, compared to GWAS *k* -mers approaches [26], that do not require themselves reference for mapping, `GraNPA` provides directly the coordinates of the regions of interest.

### Execution time

Regarding computational performance, the `GraNPA` pipeline demonstrated high efficiency across both datasets. The analysis of the *O. sativa* dataset, using a 900 MB (compressed) PVG, was completed in 2 hours and 55 minutes using 63 GB of RAM. Notably, for the cattle dataset — which involved a significantly larger 13 GB uncompressed graph — the execution time only rose to 5 hours using 232 GB of RAM. This observation suggests that `GraNPA` can handle substantial increases in data volume with a relatively limited impact on processing time, confirming its suitability for large-scale pangenomic studies.

### Limitations and Future Directions

Despite these promising results, several avenues for improvement remain. The current window-based analysis could be e.g. further refined by implementing clustering algorithms that account for both the physical position and the length of nodes, rather than a fixed node count. Moreover, while this study focused on node presence/absence, incorporating graph links (edges) as support for phenotypic information could capture more complex structural rearrangements that are currently overlooked. As the availability of high-quality *de novo* assemblies continues to grow, topology-based tools like `GraNPA` will become increasingly vital for rapid, cost-effective trait discovery in non-model species or small breeding populations.

## 5 Conclusion

We introduce here `GraNPA`, a novel framework that leverages PVG topology to directly associate genomic variations with phenotypic traits. While `GraNPA` shares functional objectives with traditional GWAS or BSA, it represents a paradigm shift in how genetic variation is handled. By operating independently on a linear reference genome, the method treats SNPs and SVs within a unified topological analysis, identifying node patterns that correlate with observed phenotypes. The efficiency of `GraNPA` was demon-strated firstly on simulated data, then across diverse biological kingdoms, providing significant results in both plant (*Oryza sativa*) and animal (*Bos tauris*) datasets, even with limited sample sizes. However, as a topology-based tool, its performance remains inherently linked to the quality of the PVG construction and the continuity of node indexing (and thus indirectly by the assembly quality of embedded haplotypes). Despite these dependencies, `GraNPA` offers a powerful, reference-free “prioritization compass” for trait discovery, paving the way for more inclusive and comprehensive genomic analyses in the pangenomic era.

## Availability and Requirements

- Project name: `GraNPA`
- Project home page: https://forge.ird.fr/diade/graphgwas/granpa
- Operating system(s): Platform independent
- Programming language: Python
- Other requirements: pysam 0.23.3, pandas 2.3.3, tqdm 4.67.1, numpy 2.4.1, statsmodels 0.14.6, matplotlib 3.10.8, PyYAML 6.0.1, scipy 1.17, GraTools 1.1
- License: GNU GPLv3
- Any restrictions to use by non-academics: *ad-hoc* license needed

### List of abbreviations

GWAS: Genome-Wide Association Study
PVG: Pangenome Variation Graph
GFA: Graphical Fragment Assembly
SNP: Single Nucleotide Polymorphism
SV: Structural Variation
PS: Phenotypic Score
BSA: Bulked Segregant Analysis
NGS: Next Generation Sequencing
RIL: Recombinant Inbreed Line
CSV: Comma Separated Value
QTL: Quantitative Trait Locus

## Declarations

### Ethics approval and consent to participate

Not applicable

### Consent for publication

Not applicable

### Availability of data and materials

The datasets generated and analyzed during the current study are available in the GranPa DataSuds Repository The rice PVG was constructed using Minigraph-Cactus using the 13 *Oryza sativa* individuals from [22], as described in [23]. All cattle chromosome graphs were originally obtained from here : https://zenodo.org/records/15097551.

### Competing interests

None

### Funding

This research was carried out within the framework of a CIFRE/ANRT PhD program, with Syngenta Seeds as the industrial partner, in collaboration with IRD and the GAIA Doctoral School.

### Code availability

Script used to generate simulated data and results are available at:

https://forge.ird.fr/diade/graphgwas/granpa and GranPa Dataverse Source Code

### Author contribution

CC prototyped, coded, validated the code and draft and finalized the manuscript. CM and FS managed the project, prototyped, validated the code, draft and finalized the manuscript.

### Supplementary information

All supplementary information are available at the GraNPA DataSuds repository.

## Acknowledgements

The authors acknowledge also the ISO 9001 certified IRD i-Trop HPC at IRD Montpellier for providing HPC resources that have contributed to the research results reported within this paper. URL: https://bioinfo.ird.fr/. We are also grateful to the GraTools team for their support and their work on the rice data, and to Nina Marthe for her collaboration on the *Sub1A* case study with GrAnnoT, as to the founders of this study, Syngenta Seeds.

## Appendix A Supplementary data

**Fig. A.1.**
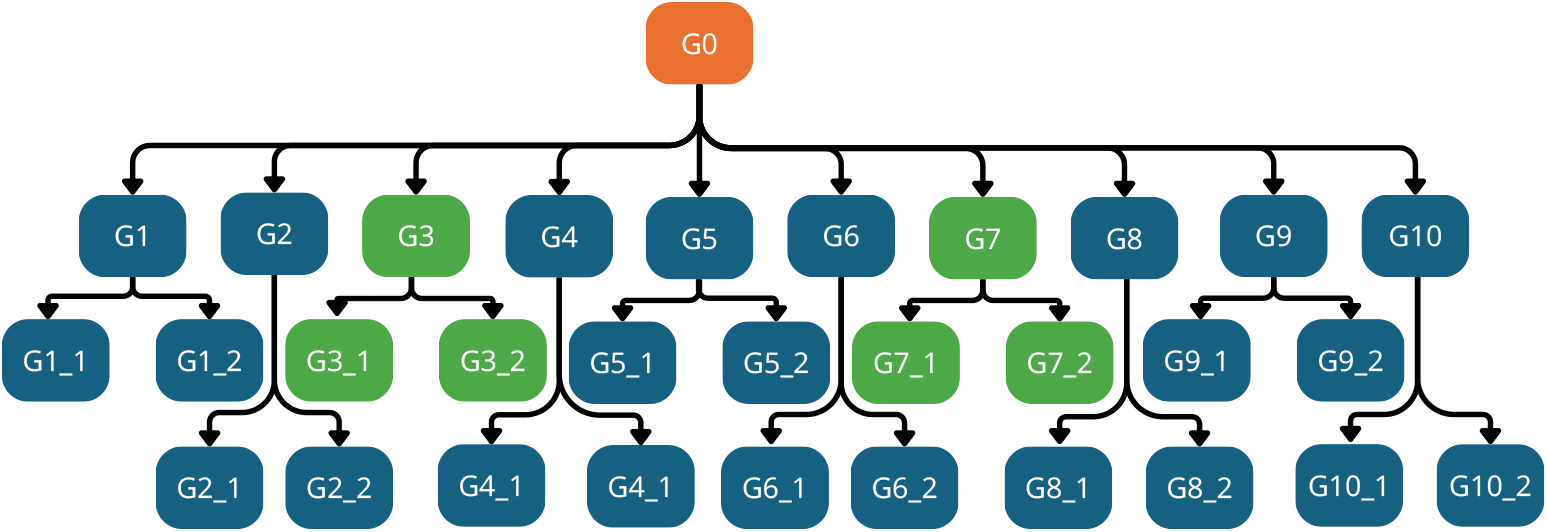
Simulated genomes data creation. The G0 genome was created by randomly generating nucleotides, each of which accounted for 25% of the 170 kb genome. Simulated genomes data creation. A 170 kb G0 genome was generated as a random nucleotide split in 4 chromosomes. A script has then been used to create variations from the G0 genome (see 2 section for more details) to generate G1 to G10. Then each G1 to G10 genomes have been used to generate 2 more genome with new variations. G0 is not included in the final PVG, and the six green genomes are used in the simulation to carry the trait. In the insertion use case, these green genomes have a 5 kb region that have been removed from all others; in the deletion use case, a 5 kb region was removed from these green genomes. The other cases shown in the appendix follow the same logic, but differ in terms of their overall structure.

**Table A.1.** Five first line of windows of interests output from both GraNPA runs on simulated data.

| **Window ID** | **Start Node** | **End Node** | **Chr** | **Mean** | **-log(p-value)** | **G3** | **G7** |
|---|---|---|---|---|---|---|---|
| **Insertion case** |  |  |  |  |  |  |  |
| 1 | 32401 | 32801 | 1 | 3.4750 | 122.44 | chr1:46182-48856 | chr1:46904-49916 |
| 2 | 32601 | 33001 | 1 | 2.9975 | 90.97 | chr1:47538-49927 | chr1:48345-50968 |
| 3 | 32201 | 32601 | 1 | 1.6775 | 32.22 | chr1:46182-47546 | chr1:46904-48353 |
| 4 | 31801 | 32201 | 1 | 1.3475 | 25.06 | chr1:44753-46182 | chr1:45520-46904 |
| 5 | 32801 | 33201 | 1 | 1.2225 | 20.86 | chr1:48856-50197 | chr1:49915-51242 |
| **Deletion case** |  |  |  |  |  |  |  |
| 1 | 31801 | 32201 | 1 | -3.0475 | 16.86 | – | – |
| 2 | 32401 | 32801 | 1 | -2.9900 | 16.86 | – | – |
| 3 | 33401 | 33801 | 1 | -2.9900 | 16.86 | – | – |
| 4 | 32801 | 33201 | 1 | -3.0475 | 16.86 | – | – |
| 5 | 33201 | 33601 | 1 | -3.0475 | 16.86 | – | – |

**Table A.2.** Samples of Asian rice in the PVG from Marthe et al [23] used in GraNPA analysis and their phenotype score.

| # | Samples | Sub species | Name | Sub1A |
|---|---|---|---|---|
| 01 | Azucena | Japonica | Azucena | -1 |
| 02 | IRGSP | Japonica | IRGSP | -1 |
| 03 | Os117425RS1 | Indica | arc | 1 |
| 04 | Os125619RS1 | Indica | LARHA MUGAD | -1 |
| 05 | Os125827RS1 | Indica | LIU XU | -1 |
| 06 | Os127518RS1 | Indica | KHAO YAI GUANG | -1 |
| 07 | Os127564RS1 | Indica | LIMA | 1 |
| 08 | Os127652RS1 | Indica | NATEL BORO | -1 |
| 09 | Os127742RS1 | Indica | PR 106 | -1 |
| 10 | Os128077RS1 | Japonica | Ketan nanka | -1 |
| 11 | Os132278RS1 | Japonica | Chao meo | -1 |
| 12 | Os132424RS1 | Indica | GOBOL SAIL (BALAM) | 1 |
| 13 | OsIR64RS1 | Indica | Ir64 | 1 |

**Table A.3.** All cattle individuals in the PVG used in GraNPA analysis and their white head phenotype.

| # | Sample | Race | White Head |
|---|---|---|---|
| 01 | ARS | Holstein | 1 |
| 02 | SIM_1 | Simmental | 1 |
| 03 | SIM_2 | Simmental | 1 |
| 04 | SIM_3 | Simmental | 1 |
| 05 | OBV_1 | Original Braunvieh | -1 |
| 06 | OBV_2 | Original Braunvieh | -1 |
| 07 | OBV_3 | Original Braunvieh | -1 |
| 08 | OBV_4 | Original Braunvieh | -1 |
| 09 | BSW_1 | Brown Swiss | -1 |
| 10 | BSW_2 | Brown Swiss | -1 |
| 11 | BSW_3 | Brown Swiss | -1 |
| 12 | BSW_4 | Brown Swiss | -1 |
| 13 | BSW_5 | Brown Swiss | -1 |
| 14 | BSW_6 | Brown Swiss | -1 |
| 15 | BSW_7 | Brown Swiss | -1 |
| 16 | BV_1 | Braunvieh | -1 |
| 17 | BV_2 | Braunvieh | -1 |
| 18 | BV_3 | Braunvieh | -1 |
| 19 | BV_4 | Braunvieh | -1 |
| 20 | BV_5 | Braunvieh | -1 |
| 21 | BV_6 | Braunvieh | -1 |
| 22 | EVO | Evolèner | -1 |
| 23 | RGV | Rätisches Grauvieh | -1 |
| 24 | HIG | Highland | -1 |

**Fig. A.2.**
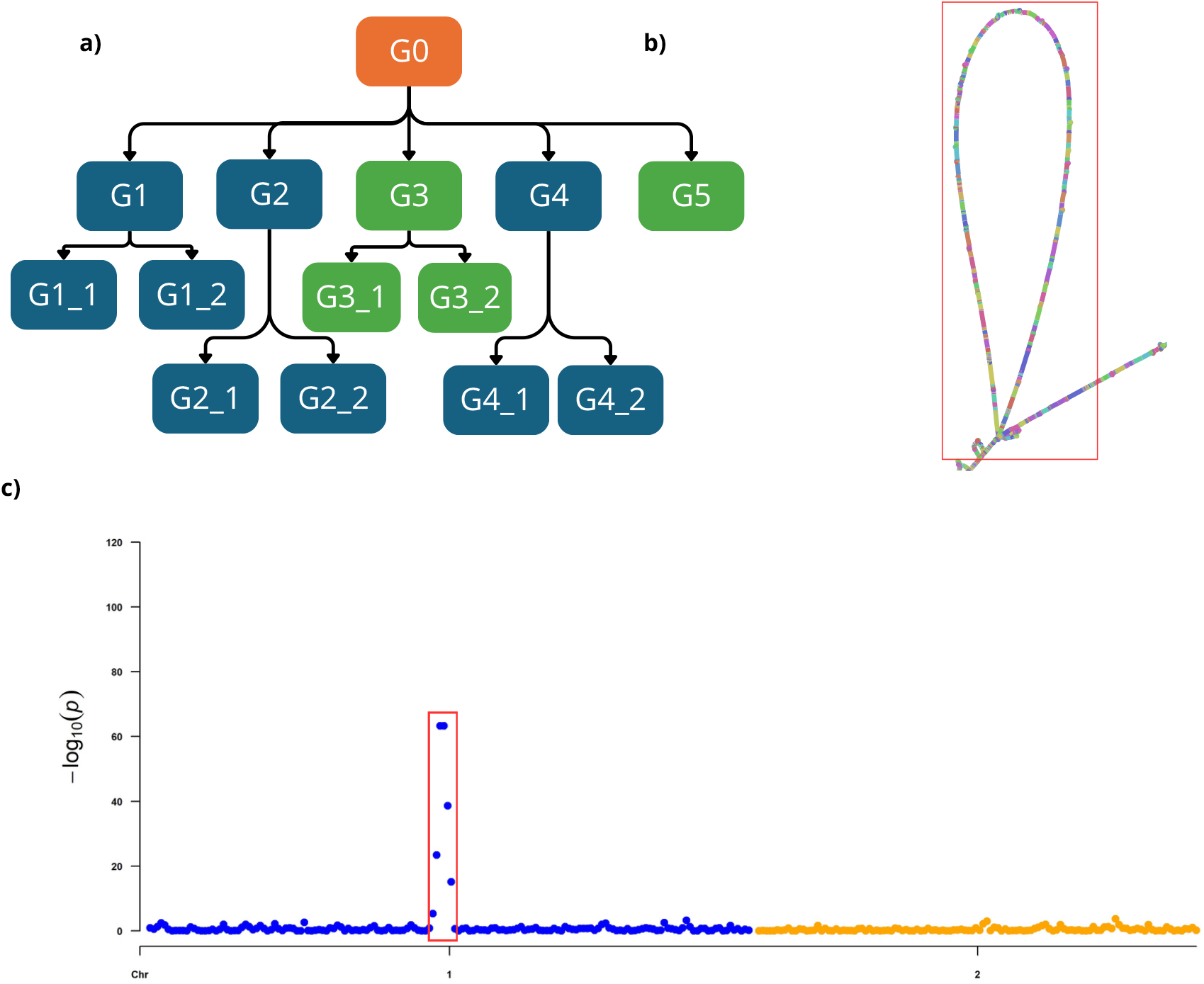
Simulated dataset with the same number of individuals and the phenotype ratio as the Asian rice data. **a)** Show the 13 individuals population structure simulated. Each arrow represents the script that simulates variations from a parent individual. Green individuals are chosen to carry a specific variation that carry the trait of interest. G0 is not used in the final graph. **b)** A bandage view of a 5 kb insertion in the individuals that carry the trait of interest. **c)** Output of a GraNPA run with the simulated graph and the encoded phenotype with 4 individuals that carry the trait at +1 and the others at -1. GraNPA was able to identify the area responsible for this simulated phenotype using a graph containing the same number of individuals and the same ratio of positive and negative phenotypes as the Asian rice dataset.

**Fig. A.3.**
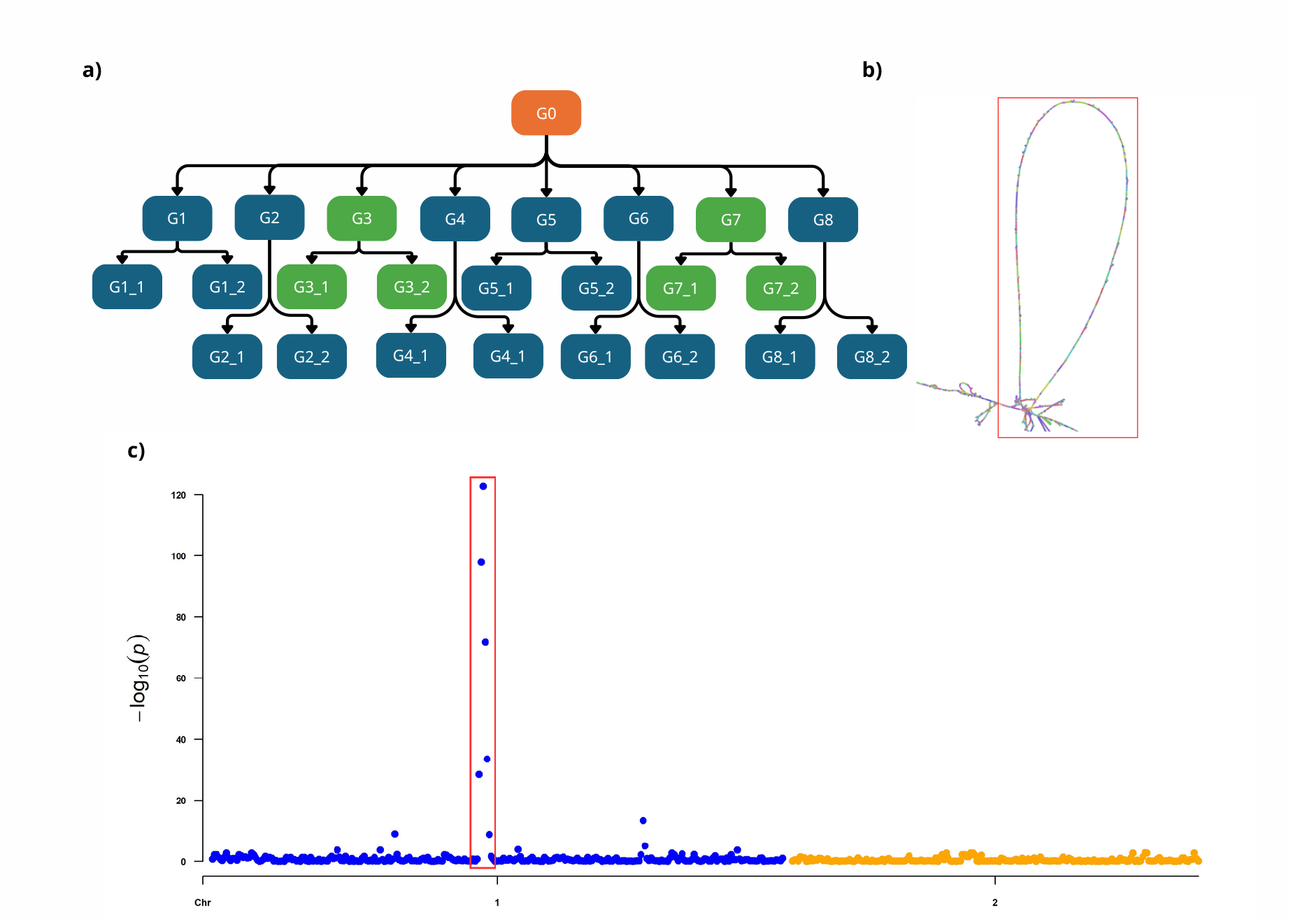
Simulated dataset with the same number of individuals and the phenotype ratio as the cattle data. **a)** Show the 24 individuals population structure simulated. Each arrow represents the script that simulates variations from a parent individual. Green individuals are chosen to carry a specific variation that carry the trait of interest. G0 is not used in the final graph. **b)** A bandage view of a 5 kb insertion in the individuals that carry the trait of interest. **c)** Output of a GraNPA run with the simulated graph and the encoded phenotype with 4 individuals that carry the trait at +1 and the others at -1. GraNPA was able to identify the area responsible for this simulated phenotype using a graph containing the same number of individuals and the same ratio of positive and negative phenotypes as the cattle dataset.

**Fig. A4.**
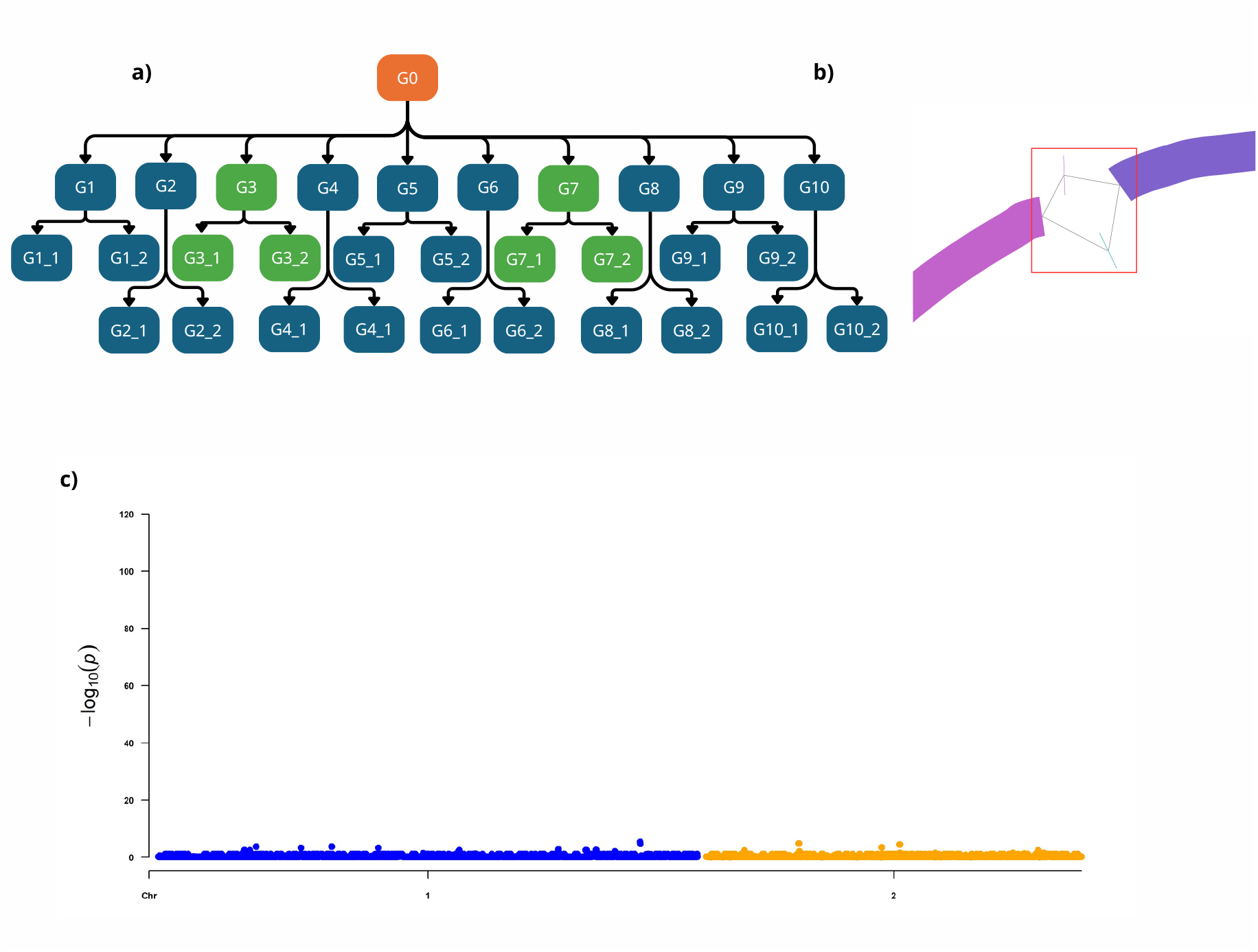
**a)** Show the 30 individuals population structure simulated. Each arrow represents the script that simulates variations from a parent individual. Green individuals are chosen to carry a specific variation that carry the trait of interest. G0 is not used in the final graph. **b)** A bandage view of the SNP, a ‘A’ in the 6 green individuals and a ‘G’ in the others. **c)** Output of a GraNPA run with the simulated graph and the encoded phenotype with 6 individuals that carry the trait at +1 and the others at -1. GraNPA was not able to identify the area responsible for this simulated phenotype (see Discussion).

**Fig. A5.**
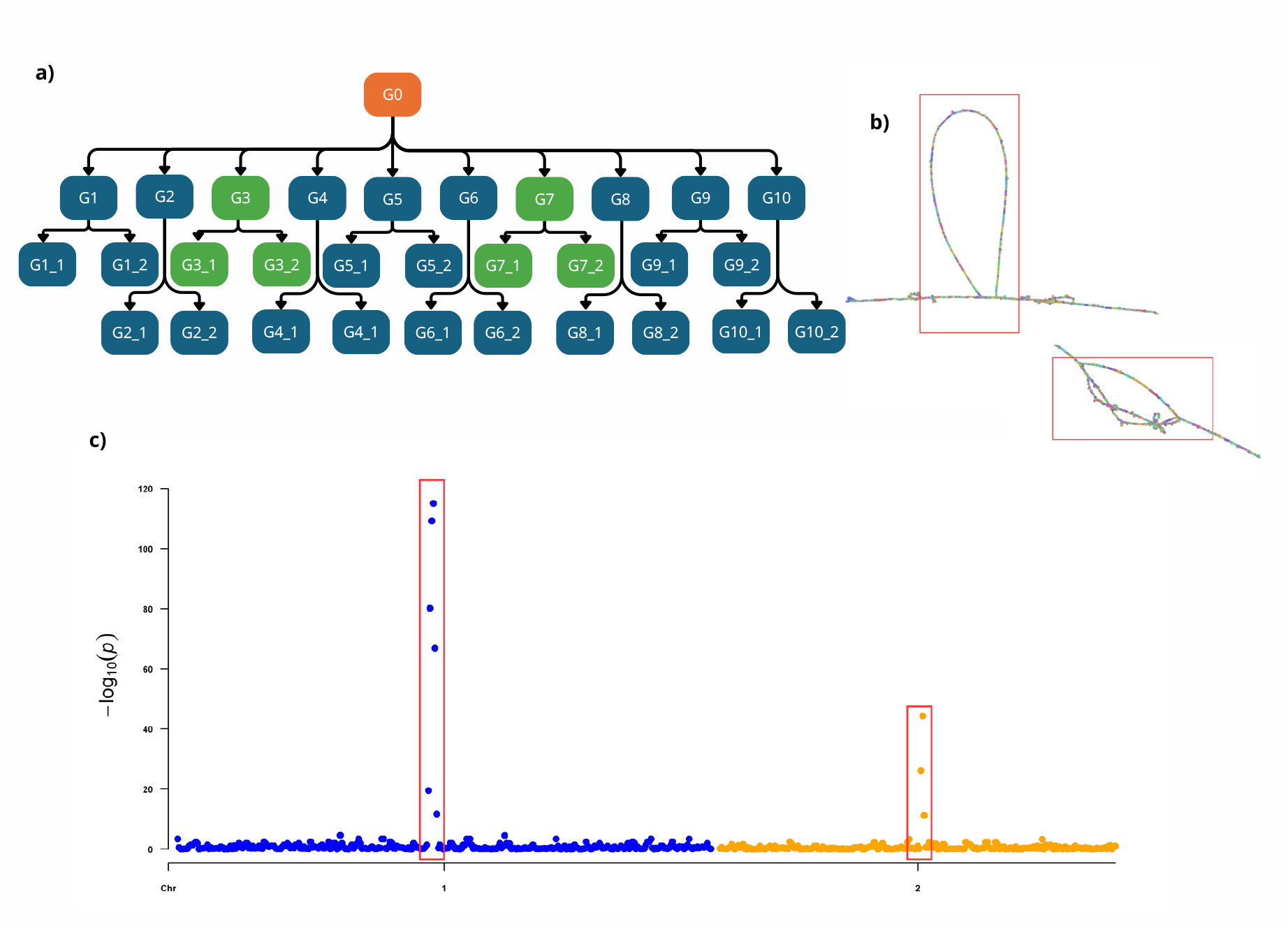
Simulated dataset with two insertion added to 6 individuals out of 30. **a)** Show the 30 individuals population structure simulated. Each arrow represents the script that simulates variations from a parent individual. Green individuals are chosen to carry a specific variation that carry the trait of interest. G0 is not used in the final graph. **b)** A bandage view of a 5 kb insertion on chromosome 1 and the 2 kb insertion in the chromosome 2 in the individuals that carry the trait of interest. **c)** Output of a GraNPA run with the simulated graph and the encoded phenotype with 6 individuals that carry the trait at +1 and the others at -1. GraNPA was able to identify both of the areas responsible for this simulated phenotype.

**Fig. A6.**
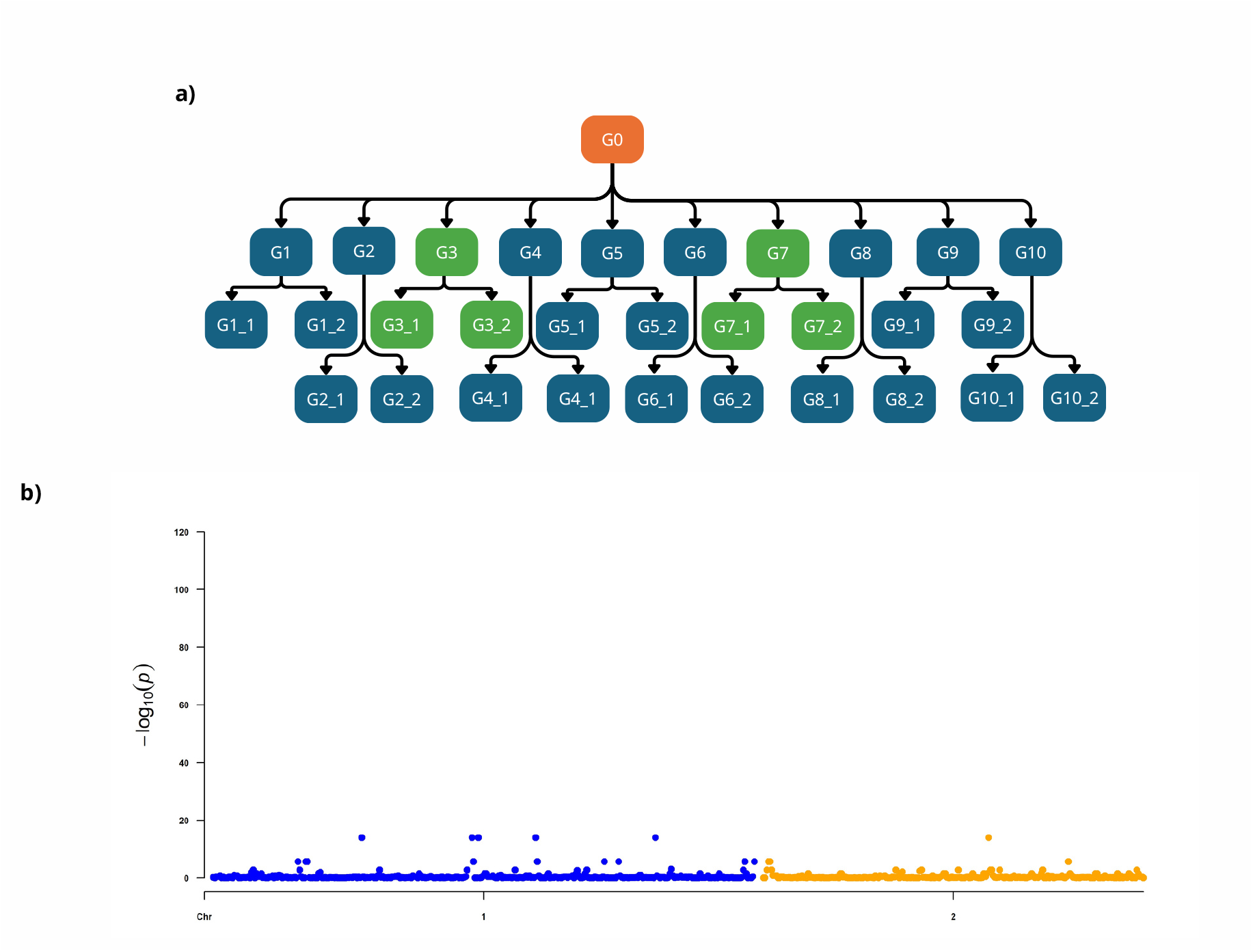
Simulated dataset with an insertion added to 6 individuals out of 30. **a)** Show the 30 individuals population structure simulated. Each arrow represents the script that simulates variations from a parent individual. Green individuals are chosen to carry a specific variation that carry the trait of interest. G0 is not used in the final graph. **b)** Output of a GraNPA run with the simulated graph and the encoded phenotype with 6 individuals that carry the trait at +1 and the others at -1. Phenotype where randomized in order to test a negative control. GraNPA did not identify any peak among the graph despite the insertion simulated in 6 individuals.

**Fig. A7.**
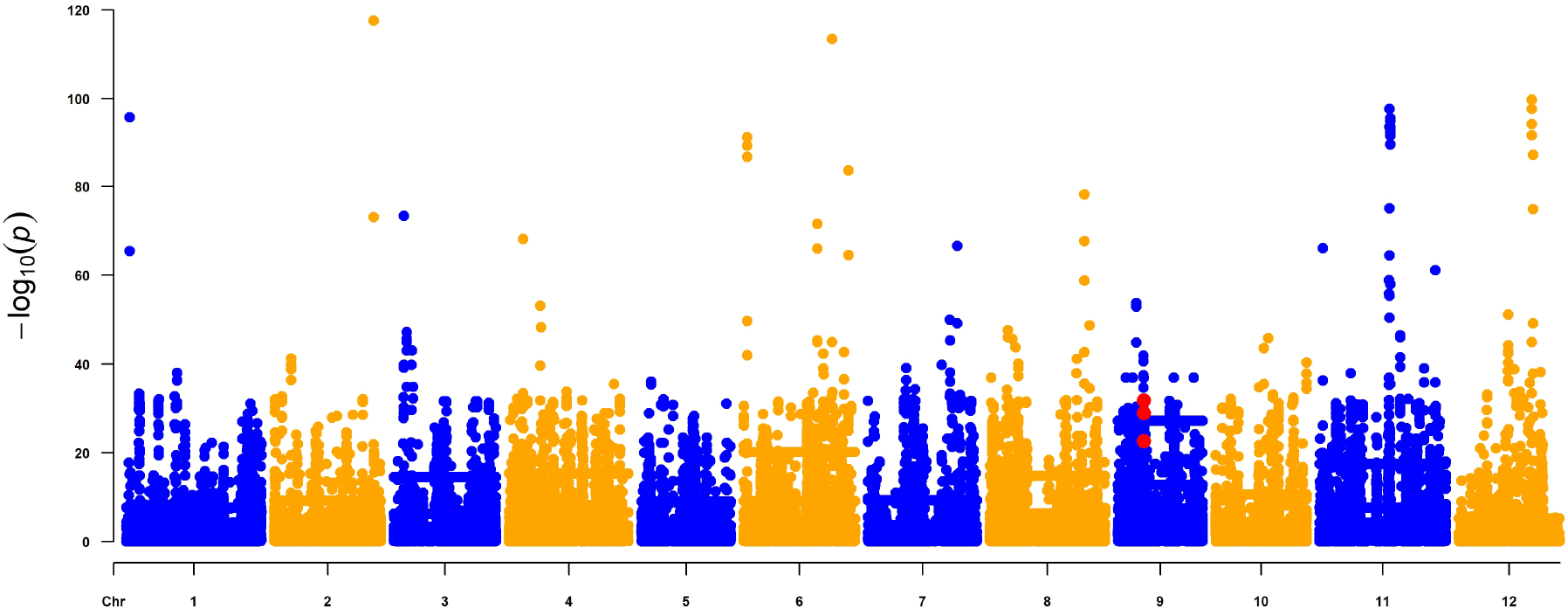
*P*-values plot of *rank compare 2indep* test on windows of 400 nodes on *Oryza sativa* PVG with the highlight (red dot) of the windows overlapping the *Sub1A-1* known position. The y-axis shows the -log of the *p*-values, and the x-axis shows the ID of the first node of each window analyzed across the (pan-)chromosomes in the PVG. The alternating colors indicate the change in chromosomes to make the plot easier to read.

## Notes

https://forge.ird.fr/diade/graphgwas/granpa

### Competing Interest Statement

The authors have declared no competing interest.

### Summary of Updates

Revisions according to reviewers after a first round of revision

